# Crop productivity of Central European Permaculture is within the range of organic and conventional agriculture

**DOI:** 10.1101/2024.09.09.611985

**Authors:** Julius Reiff, Hermann F. Jungkunst, Nicole Antes, Martin H Entling

## Abstract

Permaculture is a promising framework to design and manage sustainable food production systems based on mimicking ecosystems. However, there is still a lack of scientific evidence especially on the crop productivity of permaculture systems. In this first study on permaculture yield, we collected yield data of eleven permaculture sites in Germany and surrounding countries, that also work according to organic guidelines. We used the Land Equivalent Ratio (LER) as index to compare mixed cropping systems of permaculture sites with average monoculture yield data of the overall German agricultural sector, as well as that of only organic German agriculture. An LER of 1 indicates equal yields of the compared polyculture and monoculture, while an LER of 1.2 would indicate a 20% higher productivity of the polyculture. Mean permaculture LER as compared to total German agriculture was 0.80 ± 0.27 and 1.44 ± 0.52 as compared to German organic agriculture, both with no significant difference to 1. Our results imply, that yields of permaculture sites are comparable to predominant industrial agriculture. The observed productivity may result from the application of key permaculture principles, such as utilizing diversity and mutually supportive species and improving soil health, which contribute to system stability and resource efficiency. Provided that future studies will support our findings, permaculture could combine soil, biodiversity and climate protection with agricultural productivity. Most importantly, the variables that determine the difference in crop productivity among permaculture sites need to be identified and evaluated.

## 2 Introduction

Modern industrial agriculture, characterized by high chemical inputs, monocropping and intense soil cultivation, has led to environmental degradations such as soil erosion and loss of biodiversity (Millennium Ecosystem Assessment 2005; Foley et al. 2005; Campbell et al. 2017). While there may be a shift from southern to northern Europe and in crop types and management, no overall decline in European agricultural productivity is expected over the next few decades (Bindi and Olesen 2011). However, the frequency of extreme weather events associated with climate change leading to large-scale crop failures is increasing, e.g. in Germany (Webber et al. 2020). In response to these challenges, alternative farming approaches, that prioritize ecological sustainability and regenerative practices are gaining increased attention, such as agroecology (Barrios et al. 2020), regenerative agriculture (Schreefel et al. 2020) or diversified farming systems (Kremen et al. 2012). A promising framework for the design and management of those food production systems is permaculture (Mollison 1992; Ferguson and Lovell 2014; Krebs and Bach 2018).

Permaculture is an agroecological design system that draws inspiration from natural ecosystems and traditional and indigenous farming practices (Mollison 1992). It emphasizes the integration of a diversity of crops, with a focus on perennial and woody crops, and livestock to create self-sufficient and resilient agricultural systems (Morel et al. 2019). By mimicking the patterns and relationships found in natural ecosystems, permaculture seeks to optimize resource use, promote biodiversity and enhance ecosystem health (Ferguson and Lovell 2014). Examples for these patterns are high biodiversity, permanent soil cover, a focus on woody crops, the integration of plants and animals as well as grazing animals moving in densely packed herds (Krebs and Bach 2018). Amongst others, permaculture principles emphasize practices like polycultures, agroforestry systems, crop-livestock integration, facilitation of semi-natural habitats to enhance pest control and pollination, as well as soil conservation techniques such as mulching, composting and no-till cultivation (Reiff et al. 2024).

Implementing these principles, permaculture sites showed strong improvements in soil quality, soil carbon storage and biodiversity compared to predominant agriculture in Central Europe (Reiff et al. 2024). In addition, permaculture strives for a holistic approach that not only focuses on agricultural production but also considers social and economic aspects that aim for sustainable livelihoods and community resilience (Holmgren 2002). In addition to these improvements in sustainability, however, according to the permaculture principle ‘obtain a yield’, an agricultural system must always provide sufficient food to feed people.Although there is some evidence that permaculture can be an ecologically sustainable farming practice, there is a lack of scientific research on its crop productivity (Morel et al. 2019). The few existing studies have focused only on economic performance (Morel et al. 2015), income diversity (Ferguson and Lovell 2017) or food security based on farmer’s perception (Conrad 2014). Based on the permaculture principles outlined above we assume that crop productivity of permaculture systems is influenced by characteristics such as farm age, the size of the area under investigation, and the presence of livestock (Holmgren 2002). For instance, older farms may have had more time to establish stable and efficient ecological interactions as well as fully grown woody crops, larger investigated areas may include more diverse land uses and resources while smaller areas could be managed more efficiently and livestock integration is a key principle for nutrient cycling, pest control, and soil fertility in permaculture systems.

Therefore, this study aims to evaluate the land productivity of permaculture sites by comparing their yields to those of predominant modern agriculture in Central Europe. We used the Land Equivalent Ratio (LER) as an established tool to evaluate the productivity of mixed crop permaculture sites (Martin-Guay et al. 2018). The LER is widely used for situations with intercrops of no more than two species while evidence from combinations of three crops is scarce, with one study investigating a combination of seven crop species (Deb 2021; Deb et al. 2022). In this case, it was not feasible to conduct a single-crop experiment for every crop variety at each permaculture site. Mean values from larger samples were used to determine sole crop yields in some cases (Böhm et al. 2020), or they were estimated from the intercropping experiment itself (Seserman et al. 2018). The approach of using maximum or average sole crop yields was also described by (Mead and Willey (1980). Therefore, we used national average yield data as sole crop yield values in this study. By quantifying and comparing the yields of permaculture sites with predominant industrial agricultural systems, both overall and organic only, we provide insights into the potential benefits and limitations of adopting this approach.

## 3 Materials and methods

### 3.1 Study sites

This study evaluates yield data from eleven commercial permaculture sites in Germany (Rhineland-Palatinate, Bavaria, North Rhine-Westphalia and Lower Saxony), Switzerland, and Luxembourg, which either constitute a farm or are part of a farm. (Tab. 1). Three criteria were used for site selection. First, permaculture sites had to be designed and managed with permaculture, according to the farmer. Second, we only investigated commercial permaculture sites to focus on food production systems and to exclude permaculture sites established mainly for other purposes like subsistence or education. Third, the selected sites were required to integrate at least two different types of land use within their permaculture production systems. Examples include grazing livestock under fruit trees, or combining vegetable production with small-scale poultry farming through temporary foraging on vegetable patches and nutrient exchange. We have considered all farms in Germany and the surrounding regions, that met the specified criteria and were willing and able to provide their yield data. This data represents the agricultural production sold by the farms and was collected by the farms themselves. Yield datasets covered one year per farm between 2019 and 2022 and only crop yields from permaculture areas dedicated primarily to crop production. Livestock yields and grazing areas were excluded, as the majority of livestock production in Central Europe is based on imported forage and therefore not directly comparable in terms of land requirements. Farms were rather young with a mean age of 6 years at investigation. Therefore areas dominated by newly planted berry bushes or fruit trees, not having reached full yield potential, were excluded from the evaluation. All farms followed the principles of organic agriculture, although not all were certified. Permaculture sites 2, 3, 6 and 8 were part of a separate study on soil quality, carbon storage and biodiversity of permaculture (Reiff et al. 2024). These sites share identical identifiers in both studies.

### 3.2 Reference data

To compare permaculture yields with predominant industrial agriculture, data by the Federal Statistical Office of Germany for German agriculture of respective years was used for vegetables and strawberries (Federal Statistical Office 2023a), potatoes (Federal Statistical Office 2023b), tree fruit (Federal Statistical Office 2023c), and other soft fruit (Federal Statistical Office 2023d). These surveys are representative of Germany. As the permaculture farms in Luxembourg and Switzerland are located close to the German border, we assume that these comparative data are also representative for these farms. Data was collected from 5,100 farms in 2019 and 2020, and from 4,500 farms in 2021 and 2022 (Federal Statistical Office Germany, 2024; personal communication). Throughout Germany, most arable land parcels are used for single crop cultivation (Blickensdörfer et al. 2022). These datasets included mean crop yield data of overall (including conventional and organic) German agriculture (Y_tot_year_) and only organic German agriculture (Y_org_year_). For vegetable or fruit varieties that were not covered by these collections, mean values of respective vegetable group (such as legumes) or of all tree or soft fruit was were used for comparison (e.g. 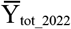(cabbage vegetables) for Y_site1_2022_(pak choi)). For organic production, vegetable yield values were only given for vegetable groups of root and tuber, fruit, leaf and stalk, cabbage and other vegetables as well as legumes (e.g. Y_org_2022_(legumes)). Thus, a ratio of organic to total agriculture was calculated for each group and year (e.g. R_2022_(legumes)=Y_org_2022_(legumes)/Y_tot_2022_(legumes)). To estimate the organic yield data of specific crop varieties, the total crop yield data of those varieties was multiplied by the respective total to organic vegetable group ratio (e.g. Y_org_2022_(sugar pea)=Y_tot_2022_(sugar pea)*R_2022_(legumes)). To estimate organic potato yield, total yield was multiplied by organic to total root and tuber vegetable ratio (Y_org_2022_(potato)=Y_tot_2022_(potato)*R_2022_(root and tuber vegetables)). For fruit tree crops organic yield data was only available for 2022, so an organic to total ratio was calculated from this data (e.g. R_2022_(apple)=Y_org_2022_(apple)/Y_tot_2022_(apple)) and applied to data of the other years (e.g. Y_org_2019_(apple)=Y_tot_2019_(apple)*R_2022_(apple). Nut crops were only grown on one permaculture site and were a relatively small proportion of total production. (Tab. 2). Nut yield data of German agriculture was not available, therefore general literature values were used for comparison of walnut (Cerović et al. 2010) and hazelnut (Erdogan 2018) yields. Tree crop organic to total ratio was applied to estimate organic nut yield values (e.g. Y_org_2022_(hazelnut)=Y_erdogan_2018_(hazelnut)*R_2022_(tree crops).

**Table 1:** Investigated Farms with permaculture. Only crop types written in italic were investigated in this study. The remaining crop types were excluded from the investigation as they were either newly planted woody crops, from areas primarily designated for livestock production, or from non-permaculture areas.

| Site | Country | Establishment | Survey | Farm area [ha] | Investigated area [ha] | Farm plant production | Farm livestock |
| --- | --- | --- | --- | --- | --- | --- | --- |
| 1 | Switzerland | 2011 | 2021 | 2,5 | 0,02 | *vegetables, soft fruit, tree crops*, grassland |  |
| 2 | Germany | 2009 | 2019 | 10 | 0,44 | *vegetables, soft fruit, tree crops*, grassland, grains | chicken, pigs, geese |
| 3 | Germany | 2009 | 2019 | 3,6 | 0,66 | *vegetables, soft fruit, tree crops*, grains | chicken |
| 4 | Switzerland | 2020 | 2021 | 5 | 0,06 | *vegetables, soft fruit, tree crops*, grassland | chicken, sheep |
| 5 | Germany | 2019 | 2021 | 1,9 | 0,22 | *vegetables, soft fruit, tree crops* | runner ducks, chicken |
| 6 | Luxembourg | 2014 | 2020 | 1,5 | 1,01 | *vegetables, soft fruit, tree crops* | runner ducks |
| 7 | Germany | 2018 | 2021 | 3,5 | 1,60 | *vegetables, tree crops* |  |
| 8 | Germany | 2013 | 2022 | 1,1 | 1,06 | *vegetables, soft fruit, tree crops* |  |
| 9 | Germany | 2022 | 2022 | 0,4 | 0,06 | *vegetables, soft fruit, tree crops*, grassland | sheep |
| 10 | Switzerland | 2015 | 2021 | 3 | 0,32 | *vegetables, soft fruit, tree crops* |  |
| 11 | Germany | 2017 | 2022 | 2,4 | 0,15 | *vegetables, soft fruit, tree crops*, grassland | chicken, pigs, sheep |

### 3.3 Land Equivalent Ratio

In all cases, permaculture sites consisted of mixed cultures of different vegetable varieties and often additional fruit trees and berry bushes. Additional integration of livestock was common, but the resulting additional animal yields could not be included in this study. The land equivalent ratio (LER) is used as an index to assess the relative productivity of these mixed crop systems compared to the mean sole crop productivity of total (both conventional and organic) and only organic German agriculture in the respective years (Mead and Willey 1980; Risch and Hansen 1982; Bomford 2009; Reynafarje et al. 2016; Paut 2018). The LER for a specific permaculture site *site* as compared to one of the management categories *man* (total or organic German agriculture) was calculated as follows

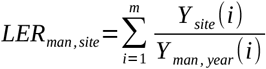

where *m* is the number of different crops yielded at the permaculture site, Y_man,year_(i) is the average monocultural yield of the *i*^*th*^ crop of respective management and year and Y_site_(i) is the yield of the *i*^*th*^ crop under intercropping of the permaculture site. Two LER values were calculated for each permaculture site, one compared to total German agriculture and one compared to German organic agriculture. An LER of 1 indicates equal productivity of the permaculture mixed system and statistical average of sole crops, while an LER of 1.2 would indicate a 20% higher productivity of the mixed system. Example calculation for yield data of permaculture site X from 2019 in comparison with total German agriculture and of just two crops:

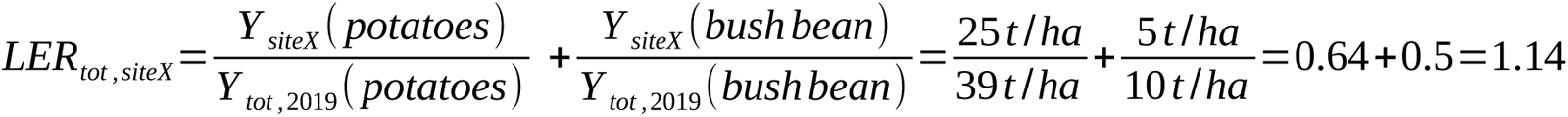

### 3.4 Statistics

Statistical analysis was carried out using R (R 4.2.1, R Development Core Team 2022). Both samples of LER values (compared to overall or organic German agriculture) were checked for normal distribution visually using the function *qqplot()* as well as mathematically using a Shapiro-Wilk-Test with the function *shapiro*.*test()*. A one sample t-Test was used to test both groups of LER values against the specified value of 1 using the function *t*.*test()*.

Two linear models were calculated using the function *lm()* with total LER or organic LER values as response variables and age, investigated area and presence of livestock at the farm level as predictor variables. Automated model selection was performed using the *dredge()* function. Model diagnostics to check for deviations from the model assumptions (normal distribution, homogeneity of variance, etc.) were performed visually using the *plot()* function on the linear model outputs. The significance of the predictor variables was evaluated with a Type II F-test using the Anova function of the ‘car’ package (Fox et al. 2023) on the full model, since no model with significant predictors was found (Table 2).

Values in the text are given as mean plus minus 0.95 confidence interval.

## 4 Results

A total of 79 crop varieties were found on the permaculture plots to calculate LER values. Of the crops considered in this study, the permaculture sites produced a total of 93.6 % vegetables, 5.8% tree crops and 0.5% soft fruit.

On average, the crop yield of permaculture sites was 21,8 ± 7,3 t ha^-1^. Table 3 displays the total crop yield and proportions of different crop types for each permaculture site. Mean permaculture site LER as compared to overall German agriculture was 0.80 ± 0.27 and 1.44 ± 0.52 as compared to only organic German agriculture (Fig. 1, Tab. 2+3). The permaculture LER of 0.80 indicates a trend suggesting that permaculture may require 20% more land to achieve the same yield as overall German agriculture. Similarly, the results indicate a trend of 44% higher permaculture productivity compared to organic German agriculture. However, both differences were not statistically significant (Tab. 2).

LER values as compared to total German agriculture and to German organic agriculture both were not significantly dependent on any of the tested predictor variables: farm age, investigated area and presence of livestock (Tab. 2).

**Figure 1:**
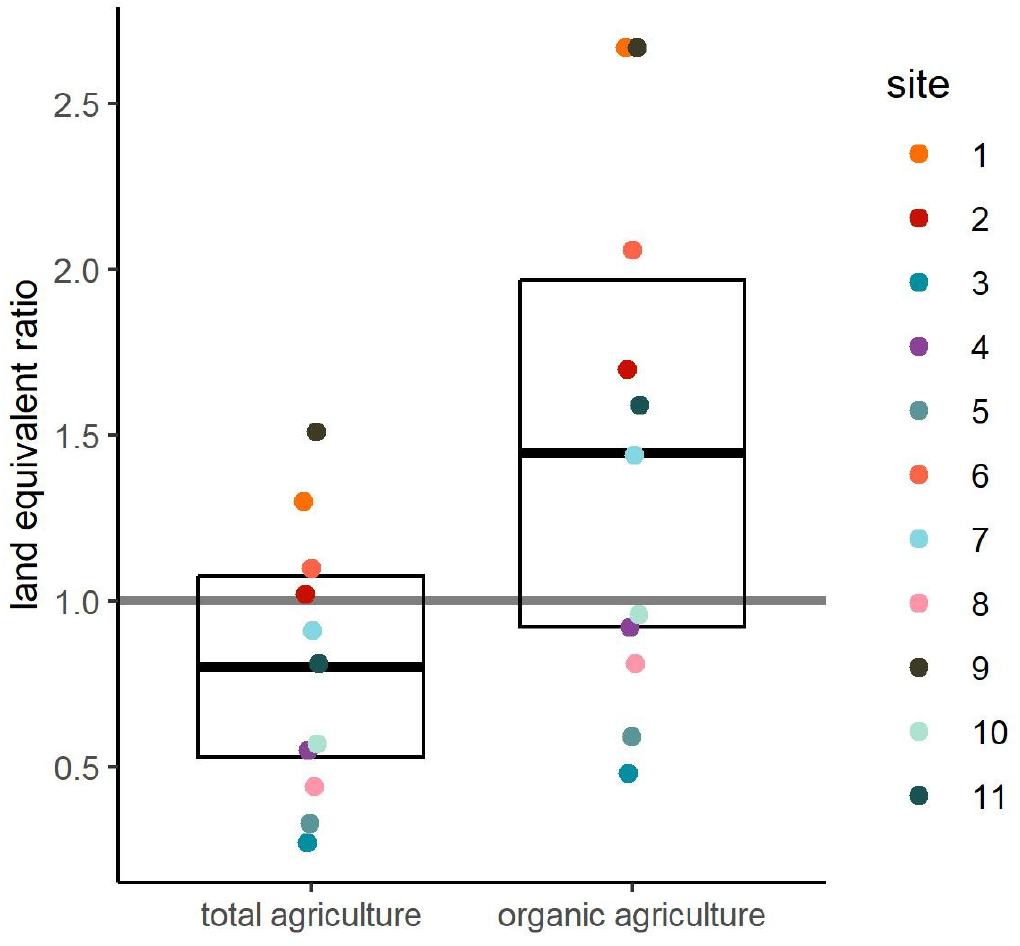
Land equivalent ratios (LER) of permaculture. LER’s of eleven permaculture sites as compared to total (p=0.137, t=-1.62, df=10) and organic (p=0.087, t=1.98, df=10) German agriculture. Bars with error bars indicate mean and 95% confidence interval, coloured dots indicate individual data points and horizontal line indicates equal land requirement of permaculture and reference.

## 5 Discussion

Both mean LER values were not significantly different from 1, indicating no significant difference in permaculture productivity compared to average German agriculture. This indicates that yields of permaculture sites are comparable to predominant industrial agriculture. With eleven farms, our sample size is relatively low, which may have limited our statistical power. Therefore, a p-value of 0.087 suggests a clear trend indicating a 44% higher productivity of permaculture farms compared to organic agriculture, although this difference is not statistically significant. This by trend higher productivity compared to German organic agriculture even suggests a potential of permaculture to bridge the productivity gap between organic and conventional agriculture, which accounts for approximately 20% lower yields in organic horticulture (Lesur-Dumoulin et al. 2017). However, as seen in Fig. 1, most farms maintained their relative position above or below an LER of 1 across both comparisons, with only a few exceptions, indicating that this potential may depend on specific farm conditions or management practices. However, LER values varied strongly between individual permaculture sites and maintained their relative position above or below an LER of 1 across both comparisons, with only a few exceptions, indicating that this potential may also depend on specific farm conditions or management practices (discussed below).. A meta study found a mean LER of 1.36 ± 0.04 with a similar range from 0.5 to 2.6 for intercropping of vegetables and/or fruit trees (Paut 2018). This value corresponds to the permaculture LER of this study as compared to German organic agriculture in general, as the permaculture farms were operated according to organic farming guidelines. As the mean permaculture LER shows a clear thrend to be substantially higher with 1.44 ± 0.52, its difference from 1 might therefore be largely explained by the use of intercropping.

The comparable yields of permaculture and industrial agriculture should also be interpreted in the context of Sustainable Development Goal 2 (Zero Hunger), which emphasizes not only the availability of sufficient food but also the importance of diverse and nutrient-rich foods to combat malnutrition (FAO 2018). Permaculture’s focus on healthy soils, achieved through practices like organic fertilisation, mulching, and minimal soil disturbance, supports the production of nutrient-dense crops (Reiff et al. 2024). Combined with the inherent crop diversity of permaculture systems, this approach has the potential to address both hunger and malnutrition while promoting sustainable agricultural practices.

It is likely, that permaculture yields are even higher than reported in this study. At some permaculture sites, yields of soft fruits, tree fruits and nuts from areas with mainly vegetable production were not recorded by the farmers. Additionally, feed provisioning from investigated areas for livestock integrated in crop production could not be taken into account in this study. Such provision constitutes an additional yield produced within the same area, reducing the need for external feeds. This includes runner ducks or chicken for permanent or temporal pest control on vegetable areas; sheep, geese or chicken grazing below woody crops or pigs fed with crops not suitable for sale.

The relatively young age of many investigated permaculture sites likely also influenced the reported yields. Permaculture systems can be understood as analogous to natural succession, where ecosystems mature over time and become increasingly stable, resilient, and productive (Shepard 2013). Similarly, a ‘mature’ permaculture system develops an intricate arrangement of mutually supporting species, which not only enhances ecological functions such as pest control and soil health but also reduces the need for external inputs like fertilizers, pesticides, and feed. This reduction in external inputs would further improve the relative productivity of permaculture systems (Holmgren 2002). In addition, higher nutrient concentrations in permaculture soils are likely to result in more nutritious food, which is essential to combat widespread malnutrition in the developing world (Reiff et al. 2024).

Beyond these biological and ecological aspects, permaculture systems also encompass a significant social dimension. They are designed to foster community involvement, knowledge sharing, and local food sovereignty (Magno 2024). These social benefits represent an additional form of ‘yield,’ although they could not be measured in this study. For example, the collaborative and participatory nature of permaculture often strengthens social networks and promotes educational opportunities within communities, which can indirectly contribute to long-term food security and resilience. While these social ‘yields’ are difficult to quantify, they are critical for understanding the broader contribution of permaculture to sustainable food systems.

**Table 2:** Statistics. Results of t-Tests and linear models on the Land-Equivalent-Ratios (LER) of 11 permaculture sites as compared to total German agriculture and to German organic agriculture fitted in R.

| Response variable | Test | Explanatory variable | t/F-value | P-value | df |
| --- | --- | --- | --- | --- | --- |
| LER (total) | One sample t-Test against 1 | NA | -1.62 | 0.137 | 10 |
| LER (total) | Linear model | Age | <0.00 | 0.995 | 7 |
| LER (total) | Linear model | Investigated area | 0.02 | 0.904 | 7 |
| LER (total) | Linear model | Presence of livestock | 0.24 | 0.641 | 7 |
| LER (organic) | One sample t-Test against 1 | NA | 1.98 | 0.087 | 10 |
| LER (organic) | Linear model | Age | 0.03 | 0.864 | 7 |
| LER (organic) | Linear model | Investigated area | 0.13 | 0.734 | 7 |
| LER (organic) | Linear model | Presence of livestock | 0.18 | 0.688 | 7 |

LER values were not significantly dependent on any of the tested predictor variables. Nevertheless, the variability of the permaculture LER values was high. Permaculture is a very context specific design tool, thus every permaculture system is different. A high variance among permaculture sites was also found for increases in soil quality, carbon storage and biodiversity compared to predominant agriculture in Central Europe (Reiff et al. 2024). We assume that variance in permaculture LER’s is a result of a combination of different factors such as the degree of complexity, the management intensity, the age of the system as well as the experience of the farmers. We did not find any dependence on the age of the permaculture sites, but our farms were also very young with a maximum age of 10 years. We suspect that this may still be a factor in older farms and in combination. The degree of complexity varied among permaculture sites and could be determined by the level of spatial and temporal integration of different land use elements. This can range from the mixed cultivation of vegetables to agroforestry and the integration of different types of livestock. A recent experiment showed, that LER’s of mixed culture of seven annual crops varied between 1.18 and 5.67 depending on cropping design (Deb 2021). Also, the level of management intensity differed between permaculture sites investigated in this study, from more extensive systems with a stronger focus on nature conservation and input efficiency to more intensive systems with a higher input of labour and resources. Ultimately, the effectiveness of a permaculture system may hinge on the farmer’s experience and competence in handling such a multifaceted system. Hence our results suggest, that well planned and managed permaculture systems are able to be as productive as prevalent industrial and especially organic agriculture. Still, on average permaculture seems to be able to reduce the yield gap of organic agriculture while still working according to its guidelines. A global meta-analysis revealed that, mean organic agriculture yields were 25% lower compared to those of conventional agriculture (Seufert et al. 2012). At the same time, permaculture seems to strongly improve environmental conditions of the agroecosystem in terms of soil quality, carbon storage and biodiversity (Reiff et al. 2024).

**Table 3:** Crop yield of permaculture sites. Land-Equivalent-Ratio of eleven permaculture sites in Germany and neighbouring countries as compared to overall (LER total) and only organic (LER organic) German agriculture. Yield includes crop yield of vegetables, tree crops and soft fruit. The proportions of vegetable groups, soft fruit, tree fruit and tree nut in the total yield of the permaculture site are given as percentage values.

| site | LER total | LER organic | yield [t/ha] | root/tuber veg. [%] | fruit veg. [%] | cabbage veg. [%] | leaf/stalk veg. [%] | legume [%] | other veg. [%] | soft fruit [%] | tree fruit [%] | tree nut [%] |
| --- | --- | --- | --- | --- | --- | --- | --- | --- | --- | --- | --- | --- |
| 1 | 1,30 | 2,67 | 20 | 4 | 68 | 1 | 13 | 0,5 | 0,0 | 13,4 | 0,0 | 0,0 |
| 2 | 1,02 | 1,70 | 17 | 30 | 18 | 21 | 26 | 4,8 | 0,0 | 0,0 | 0,0 | 0,0 |
| 3 | 0,27 | 0,48 | 32 | 29 | 33 | 14 | 7 | 2,5 | 0,0 | 1,4 | 11,8 | 0,3 |
| 4 | 0,55 | 0,92 | 7 | 37 | 37 | 6 | 18 | 0,5 | 0,0 | 2,1 | 0,0 | 0,0 |
| 5 | 0,33 | 0,59 | 31 | 21 | 24 | 17 | 20 | 1,4 | 0,0 | 0,1 | 17,0 | 0,0 |
| 6 | 1,10 | 2,06 | 12 | 17 | 39 | 10 | 29 | 4,2 | 0,1 | 0,0 | 0,0 | 0,0 |
| 7 | 0,91 | 1,44 | 7 | 27 | 25 | 3 | 41 | 3,9 | 0,0 | 0,0 | 0,0 | 0,0 |
| 8 | 0,44 | 0,81 | 32 | 37 | 21 | 27 | 15 | 0,6 | 0,0 | 0,0 | 0,0 | 0,0 |
| 9 | 1,51 | 2,67 | 45 | 27 | 41 | 13 | 14 | 4,0 | 0,0 | 0,2 | 0,0 | 0,0 |
| 10 | 0,57 | 0,96 | 11 | 19 | 9 | 17 | 31 | 6,0 | 0,1 | 6,4 | 11,4 | 0,0 |
| 11 | 0,81 | 1,59 | 26 | 13 | 33 | 7 | 44 | 0,3 | 2,1 | 0,0 | 0,0 | 0,0 |

Common permaculture literature suggests to rely on annual crops until woody crops are established and reaching full yield (Shepard 2013; Perkins 2016). The high contribution of vegetables to the farms’ total production found on all permaculture sites in this study aligns with their recent establishment (Tab 1, Tab 3). The viability of permaculture sites relying mainly on vegetables could be evidenced in a case study in France. Here, on a permaculture site measuring 1000 m^2^ one person produced an income ranging from 900 to 1600 € per month, with a mean workload of 43 hours per week (Morel et al. 2015). In addition, a study in the USA found permaculture farms to fit well within the emerging framework of diversified farming systems, with a high diversity of production and income, including non-production enterprises, to develop and maintain diverse agroecosystems (Ferguson and Lovell 2017). In Malawi, farmers experienced economic and nutritional benefits from utilizing permaculture through increased, more diverse and more stable yields (Conrad 2014). This first study on permaculture yields in Central Europe demonstrates that permaculture also has the potential to compete with industrial methods in temperate climates. This first study on permaculture yields in Central Europe demonstrates that, at the farm level, permaculture has the potential to achieve productivity comparable to industrial methods in temperate climates. However, the scalability of permaculture systems to larger surfaces remains a challenge that warrants further research.

## 6 Conclusion

Our findings suggest that well-planned and managed permaculture systems can obtain productivity levels comparable to industrial agriculture while adhering to guidelines of organic agriculture. This highlights the potential of permaculture to bridge the productivity gap between organic and conventional agriculture, while regenerating agroecosystems. Further promotion and adoption of permaculture principles could enhance sustainable food production and reduce reliance on industrial farming methods.

The limited scope of this study with eleven sites and yield data from only one year needs further and larger studies to confirm our results. In addition, the high variance of LER values among individual permaculture sites indicates the need for more research focused on understanding the factors influencing productivity in permaculture systems. Future studies should investigate larger samples of permaculture systems from different continents and climates, as well as compare factors such as system complexity (e.g., number of integrated crops and livestock, management intensity (e.g., labor or inputs per hectare), and farmer experience (e.g., years of permaculture practice or training) to determine their impact on permaculture yields. Additionally, exploring long-term effects of older permaculture systems, including staple crop (e.g. grains) and livestock yield, and comparing them to conventional agricultural practices would provide valuable and much needed insights.

## 7 Acknowledgments

We thank the Heinrich-Böll-Foundation for funding a PhD scholarship supporting this research and all farmers involved for making this study possible.

## 9 Availability of data and material

The datasets generated during and/or analyzed during the current study will be made openly available.

## 11 Author Contributions

Funding acquisition, methodology development and original draft preparation were done by Julius Reiff. Data acquisition and analysis was done by Julius Reiff and Nicole Antes. Conceptualization was done by Hermann F Jungkunst, Martin H Entling and Julius Reiff. Review and editing was done by all Co-Autors.

